# Resistance to protoporphyrinogen oxidase inhibitor herbicides in giant ragweed (*Ambrosia trifida*) is associated with a novel R98Q target-site mutation in PPO2

**DOI:** 10.64898/2026.07.16.738972

**Authors:** Cristiana Bernardi Rankrape, Isabel Werle Noe, Eduardo Lago, Alexander J. Lopez, Patrick J. Tranel, Karla L. Gage

## Abstract

**BACKGROUND:** Reduced control of giant ragweed (*Ambrosia trifida* L.) with protoporphyrinogen oxidase (PPO)-inhibiting herbicides was recently reported in two southern Illinois populations, VRC and TMS. The objectives of this study were to assess resistance to postemergence-applied PPO inhibitors in VRC and TMS, evaluate VRC response to acetolactate synthase (ALS)- and enolpyruvyl shikimate phosphate synthase (EPSPS)-inhibiting herbicides, and identify target-site mechanisms associated with PPO-inhibitor resistance.

**RESULTS:** Based on LD estimates, VRC resistance ratios ranged from 1.1- to 3.7-fold for lactofen and 2.2- to 6.9-fold for fomesafen relative to PPO-sensitive populations SIU and DSO. In TMS, LD estimates were 257.4 g ai ha ¹ for lactofen and 318.4 g ai ha ¹ for fomesafen. Glyphosate LD estimates exceeded three times the labeled field rate in VRC, SIU, and DSO, whereas VRC and SIU had higher cloransulam-methyl LD estimates than DSO. Whole-transcriptome sequencing identified polymorphisms in *PPX1* and *PPX2*; however, only PPO2 R98Q altered a catalytic-domain binding-pocket residue and was considered likely to contribute to resistance in VRC. R98Q was absent in TMS, and *PPX1* and *PPX2* expression did not differ among populations.

**CONCLUSION:** VRC and TMS have evolved resistance to lactofen and fomesafen, and VRC also exhibited reduced sensitivity to cloransulam-methyl and glyphosate. PPO2 R98Q is novel in *A. trifida* and, to our knowledge, represents the first report of this mutation associated with PPO-inhibitor resistance in plants. The absence of target-site alterations in TMS suggests a potential non-target-site basis for resistance. These findings highlight the need for integrated, diversified management to further reduce herbicide selection pressure.

## 1 INTRODUCTION

Giant ragweed (*Ambrosia trifida* L.) is a summer annual broadleaf weed native to North America and was ranked as the third most troublesome weed in soybean [*Glycine max* (L.) Merr.] production systems in the 2025 survey of weeds in broadleaf crops, behind only waterhemp [*Amaranthus tuberculatus* (Moq.) Sauer] and Palmer amaranth (*Amaranthus palmeri* S. Watson) (1, 2). Although *A. trifida* has spread to nearly 40 countries across multiple continents, it is considered invasive in only a small proportion of these regions. Nevertheless, its widespread distribution and highly competitive ability have made it a major agricultural concern (3, 4).

The competitive ability of *A. trifida* is largely attributed to several advantageous biological characteristics, including germination across a broad range of temperatures and soil depths, and early-season emergence (2, 5, 6). Once emerged, *A. trifida* plants exhibit rapid vertical growth and a highly leafy growth habit that enables them to maintain dominance within crop canopies, causing substantial yield losses even at low densities. For instance, soybean yield reductions of up to 77% have been reported with densities as low as one *A. trifida* plant m ², while corn yield losses of approximately 61% occur at densities of 13.8 plants 10 m^-^² (5, 7, 8). In addition to its weediness, *A. trifida* also poses significant human health risks due to the production of highly allergenic pollen associated with respiratory and dermatological illness (9, 10).

In recent years, management of *A. trifida* has become more challenging due to the evolution of herbicide resistance in the species. To date, *A. trifida* populations have evolved resistance to acetolactate synthase (ALS)-inhibitors, enolpyruvyl shikimate phosphate synthase (EPSPS)-inhibitor, and, more recently, to protoporphyrinogen oxidase (PPO)-inhibitors. All documented cases of herbicide-resistant *A. trifida* populations originate from the United States (U.S.) and Canada. Within the U.S., most resistance cases have been reported in the midwestern region (11–13).

PPO-inhibiting herbicides are commonly used for preemergence and postemergence control of broadleaf weed species, with limited activity on grasses particularly due to plant morphology (e.g., leaf angle) and herbicide uptake barriers (14). The herbicidal activity of these herbicides results from the disruption of the tetrapyrrole biosynthesis pathway by the inhibition of PPO enzymes responsible for catalyzing the conversion of protoporphyrinogen IX (protogen) to protoporphyrin IX (proto) needed for chlorophyll and heme production. Consequently, the substrate protogen accumulates and leaks into the cytoplasm, where it is spontaneously oxidized to a photoreactive variant of proto. In the presence of light and oxygen, this leads to the formation of singlet oxygen and other reactive oxygen species, resulting in loss of membrane integrity and cellular collapse through lipid peroxidation (15–17).

All land plants are known to possess two distinct PPO isoforms derived from an ancient gene duplication that differ in subplastidial localization: PPO1 and PPO2 (18). The PPO1 isoform is localized to the chloroplast thylakoid membranes, while the PPO2 isoform is localized primarily to the chloroplast envelope. PPO2 was previously believed to be localized to the mitochondria based on early studies with tobacco (*Nicotiana tabacum* L.) but has more recently been shown to be localized exclusively to the chloroplast in *Arabidopsis thaliana* (19–21). However, PPO2 localization patterns may vary among species at the subplastidial level, as evidence from Amaranthaceae indicates that PPO2 can localize to both chloroplast envelope and thylakoid membranes (21).

To date, at least 18 weed species have evolved resistance to PPO-inhibitors globally, (12), with most cases linked to target-site (TS) mutations in *PPX2* that result in amino acid substitutions or deletions in PPO2, often affecting residues in or near the catalytic domain (14, 22). Recently, PPO-inhibitor resistance was confirmed in a Wisconsin *A. trifida* population and was attributed to a mutation in *PPX2* resulting in an arginine to leucine substitution at position 98 (R98L) in PPO2, a TS mutation previously documented in *A. artemisiifolia* (11, 23).

Although several PPO-inhibiting herbicides can still provide high levels of control (>95%) in most *A. trifida* populations, continued reliance on these chemistries is likely to foster the evolution of resistant biotypes (24–26). Indeed, a farmer with a history of PPO-inhibiting herbicide use recently reported reduced efficacy of this class of herbicides in the southern Illinois (IL) field populations designated VRC and TMS. Therefore, the objectives of this study were to assess resistance to postemergence-applied PPO inhibitors in the VRC and TMS populations, evaluate the response of VRC to ALS- and EPSPS-inhibiting herbicides, and determine whether TS-based mechanisms were associated with reduced efficacy of PPO inhibitors in these populations.

## 2 MATERIALS AND METHODS

### 2.1 Plant materials and dose-response experiments

Seeds of the suspected PPO-resistant populations, VRC and TMS, were collected in the fall of 2024 from fields located in Alexander County, IL, located approximately 8 km apart, and managed by the same farmer. Seeds were a composite sample of approximately 40 plants, a sample size recommended for outcrossing species such as *A. trifida* (27, 28). Two PPO-sensitive populations, SIU and DSO, were included for characterizing resistance in the greenhouse screenings. These two populations were collected from different fields located approximately 18 km apart in Jackson County, IL, and preliminary screenings indicated that these populations had a 90% control with the PPO-inhibitors lactofen and fomesafen (data not shown).

Seeds from all four populations were initially hand-threshed and cleaned using an aspirator and subsequently placed in a 45.4 x 19.6 x 19.6 cm delicate mesh bag, buried in water-saturated sand, and kept outdoors, exposed to natural environmental conditions for approximately 10 weeks (November 18^th^, 2024 to January 25^th^, 2025) to break dormancy. After this period, seeds were sieved to separate them from the sand, placed in a plastic bag, and stored at 4 °C until further use. Subsequently, dose-response experiments were conducted at the Southern Illinois University Horticultural Research Center (HRC), organized in a completely randomized design, with eight replications per treatment, and repeated once over time. Seeds from each population were first sown into 164-cm³ containers containing a custom potting mix consisting of approximately 75–85% peat moss organic matter and 10–20% perlite between a pH range of 4.5–6.0. Individual seedlings with the first true leaves were transplanted into 0.7-L plastic pots filled with the same potting mix described previously. After the transplant, plants were watered daily to maintain soil moisture and fertilized every three days with 20-20-20 water-soluble fertilizer (Peters Professional^®^ Soluble Plant Food, Scotts-Sierra Horticultural Products Co., Marysville, OH 43041) delivering 200 ppm of nitrogen (N), phosphorus (P), and potassium (K).

Greenhouse settings were set to maintain a 16-hour photoperiod, and light was supplemented with 1000-W high-pressure sodium (HPS) fixtures. Mean day and night temperatures were 29 °C and 24 °C, respectively, with relative humidity averaging 28% during the day and 35% at night. Herbicides were applied when plants reached 5 to 10 cm in height using a spray chamber (DeVries Manufacturing Inc, Hollandale, MN, USA) equipped with an 8002EVS nozzle (TeeJet^®^ Technologies, Glendale Heights, IL), calibrated to deliver 140 L ha ¹ at 207 kPa. Herbicide rates used were 0, 0.03, 0.5, 1, 2, and 4x the label rate of fomesafen [Flexstar^®^; Syngenta Crop Protection, LLC, Greensboro, NC, USA; 1×: 263 g active ingredient (a.i.) ha^−1^], lactofen [Cobra^®^; Valent Corporation, San Ramon, CA, USA; 1×: 220 g a.i. ha^−1^], cloransulam-methyl [FirstRate^®^; Corteva Agriscience, Indianapolis, IN, USA; 1×: 18 g a.i. ha^−1^], and glyphosate [Roundup Powermax ^®^; Bayer Crop Science, St Louis, MO, USA; 1×: 1260 g acid equivalent (a.e.) ha^−1^]. The adjuvant ammonium sulfate (AMS) at 2.5% v/v was used for all herbicides and crop oil concentrate (COC) at 1% v/v was used for all except glyphosate (Table 1). At 21 days after application (DAA) plant survival was recorded as 0 (dead) or 1 (alive).

**Table 1.** Description of herbicide materials used to evaluate resistance in the *Ambrosia trifida* populations in this study.

| Herbicide | Trade name | WSSA SOA<sup>†</sup> | Application rate (g ai ha<sup>-1</sup> or ae ha<sup>-1</sup>)<sup>‡</sup> | Adjuvant (v/v %)<sup>§</sup> | Manufacturer |
| --- | --- | --- | --- | --- | --- |
| Cloransulam-methyl | FirstRate<sup>®</sup> | ALS (2) | 18 | COC (1) + AMS (2.5) | Corteva Agriscience, Indianapolis, IN |
| Fomesafen | Flexstar<sup>®</sup> | PPO (14) | 263 | COC (1) + AMS (2.5) | Syngenta Crop Protection, Greensboro, NC |
| Lactofen | Cobra<sup>®</sup> | PPO (14) | 220 | COC (1) + AMS (2.5) | Valent U.S.A. LLC, San Ramon, CA |
| Glyphosate | Roundup Powermax<sup>®</sup> | EPSPS (9) | 1260 | AMS (2.5) | Bayer CropScience, St. Louis, MO |

<sup>†</sup> Abbreviations: Weed Science Society of America (WSSA) Herbicide Site of Action (SOA): ALS, acetolactate synthase (Group 2); EPSPS, enolpyruvyl shikimate phosphate synthase (Group 9); PPO, protoporphyrinogen oxidase (Group 14).
<sup>‡</sup> The application rate represents the 1x rate based on the respective herbicide label, and recommendations for controlling *Ambrosia trifida* when specified. ai, active ingredient, ae, acid equivalent.
<sup>§</sup> Abbreviations: COC, crop oil concentrate; AMS, ammonium sulfate.

To model the responses of *A. trifida* populations to each herbicide, plant survival data were fit to a two-parameter log-logistic model for each herbicide separately (29) using the *drc* package version 3.0-1 (30) in *R* version 4.6.0 (31) and *RStudio* version 2025.10.31.452 (32). The two-parameter log-logistic model was (Equation 1):

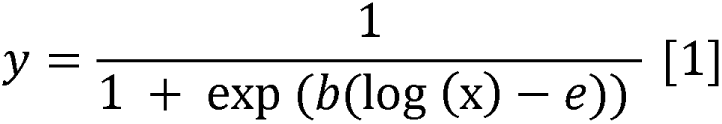

where *y* is the response variable (survival), *x* is the herbicide rate, *b* is the slope around the inflection point, and *e* is the herbicide rate causing 50% reduction in survival (LD_50_).

### 2.3 Transcriptome sample preparation and sequencing

A whole-transcriptome sequencing approach was conducted to assess potential TS resistance mechanisms associated with PPO-inhibitor resistance in the VRC and TMS populations. Plants from the two PPO-resistant populations, VRC and TMS, and a PPO-sensitive population, SIU, were grown under the same greenhouse conditions described previously. Leaf tissue samples were collected prior to herbicide treatments when plants reached 5 to 10 cm in height and stored at −80 °C for subsequent RNA extraction. Following sample collection, plants were treated at the 1× rate and 2× rate of fomesafen [Flexstar; Syngenta Crop Protection; 1× = 396 g a.i. ha^−1^ + 1% v/v COC + 2.5% v/v AMS] and lactofen [Cobra; Valent Corporation; 1× = 218 g a.i. ha^−1^ + 1% v/v COC + 2.5% v/v AMS]. Following herbicide treatments, two samples from the sensitive population and five samples from resistant populations were retained for sequencing. Detailed information for all samples is provided in Supplementary Table 1.

Total RNA was extracted using a TRIzol-based method (33), followed by DNase I treatment (Invitrogen, Life Technologies Corporation, Carlsbad, CA, USA) to remove contaminating DNA. Prior to sequencing, RNA quality and quantity were assessed through gel electrophoresis on a 1% agarose gel and with a Qubit 4.0 Fluorometer (Thermo Fisher Scientific, Waltham, MA, USA), respectively. RNA samples were then sent to the Roy J. Carver Biotechnology Center at the University of Illinois for Illumina library preparation and sequencing on an Illumina NovaSeq X 25B lane to generate 150-bp paired-end reads. Raw sequencing reads have been deposited in the NCBI Sequence Read Archive (10) under BioProject accession PRJNA1470601, with SRA accession numbers SRR38842042–SRR38842048.

### 2.4 Screening for TSR-based mechanisms in *PPX1* and *PPX2*

Sequenced read quality was assessed using FastQC v0.12.0 (34) and summarized with MultiQC v1.12 (35), after which reads were aligned to an *A.* trifida genome AAC_Ahel_1.0.Atri_hap2 available on NCBI (Genbank accession GCA_030844895.1) using STAR v2.0.6 (36, 37). Subsequently, variant calling was performed on the aligned reads using BCFtools v1.20 (38). Joint variant calling was restricted to the genomic intervals corresponding to *PPX1* and *PPX2*. For each region, genotype likelihoods were generated with BCFtools using a maximum read depth of 10,000, a minimum mapping quality of 20, and a minimum base quality score of 20. Coding variant effects were assigned using a custom script in Python v3.10.12, and variants were classified as synonymous or missense.

To identify missense substitutions that may be contributing to resistance in these populations, translated consensus sequences of the PPO1 and PPO2 proteins were first compared with multiple species to evaluate the conservation at these sites (Supplementary Table S2). Multiple sequence alignments using the consensus protein sequences were performed against sequences from both closely related and distantly related species obtained from NCBI using CLC Sequence Viewer v8.0 (CLC Bio, Aarhus, Denmark). Furthermore, missense substitutions were evaluated for their potential to influence herbicide binding by examining their localization relative to the substrate binding pocket in the catalytic domain of the PPO2 crystalized structure (1SEZ) from tobacco (39) using UCSF ChimeraX v1.8 (40).

To assess differences in *PPX* gene expression across *A. trifida* populations, read counts were obtained from aligned reads using featureCounts v2.0.6 (41) from the Subread package. Trimmed mean of M-values (TMM) normalization factors were estimated from the raw read count matrix using edgeR v4.0.16 (42). *PPX1* and *PPX2* expression values were then calculated as TMM-normalized log counts per million (logCPM) using gene-specific counts and TMM-adjusted library sizes.

## 3 RESULTS AND DISCUSSION

### 3.1 *Ambrosia trifida* responses to PPO-, EPSPS-, and ALS-inhibitors

Greenhouse dose-response experiments indicated that responses varied among herbicides and sites of action across *A. trifida* populations. Responses of VRC and TMS populations were compared with those of two PPO-sensitive populations, SIU and DSO. For the PPO-inhibiting herbicide lactofen, estimated LD_50_ (+/- SE) values for VRC, SIU, and DSO were 87.1 (±19.6), 23.5 (±8.8), and 82.5 (±91.4) g a.i ha^-1^, respectively, resulting in resistance ratios for VRC ranging from 1.1- to 3.7-fold relative to SIU and DSO (Table 2; Figure 1A). Although DSO had an estimated LD_50_ similar to VRC, the large standard error associated with this estimate suggests that this response should be interpreted cautiously. For fomesafen, estimated LD_50_ values for the VRC, SIU, and DSO were 74.5 (±23.6), 33.1 (±124.2) and 10.8 (±9.1) g a.i ha^-1^, respectively, indicating resistance ratios ranging from 2.2- to 6.9-fold (Table 2; Figure 1B).

**Figure 1.**
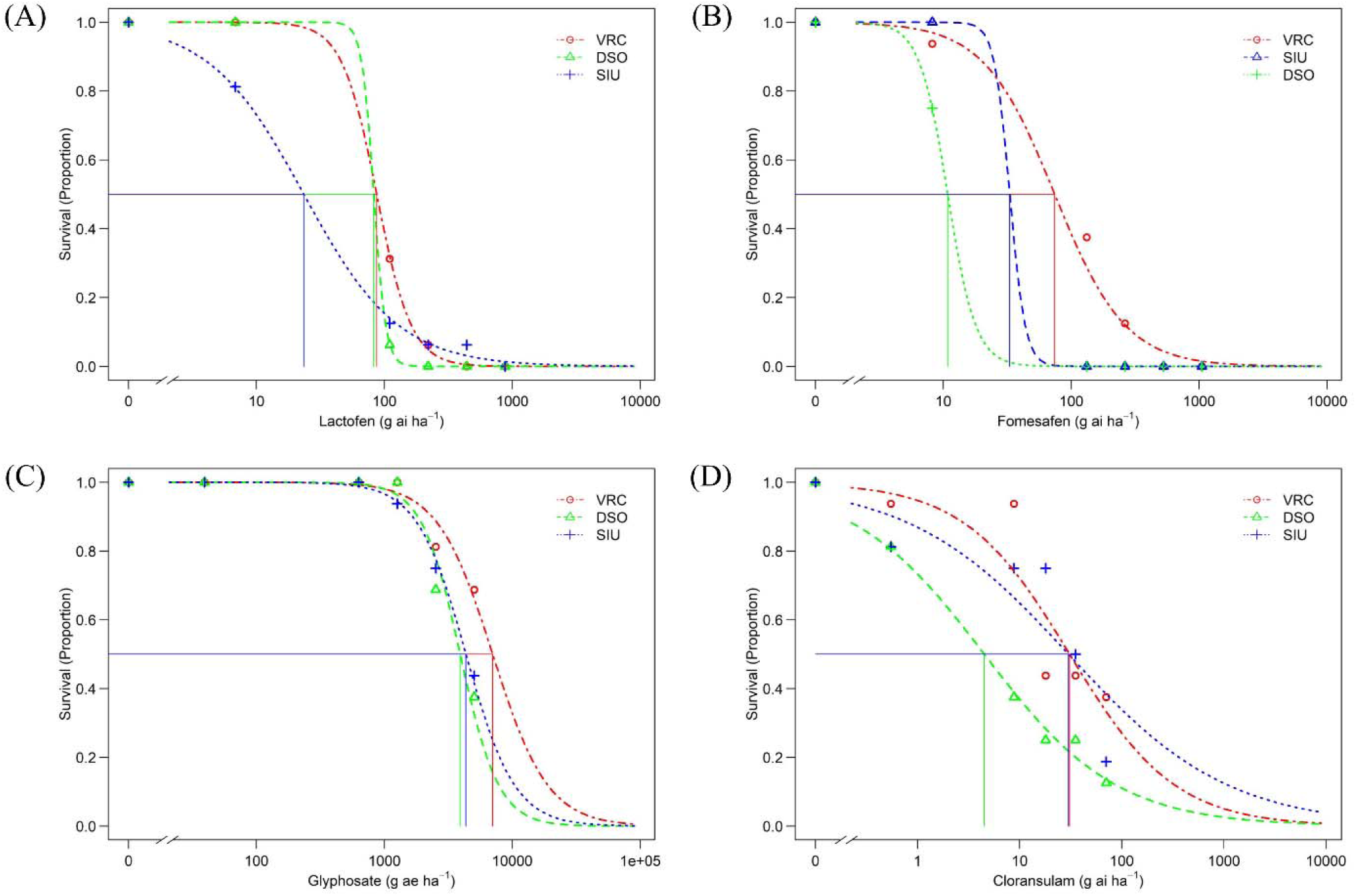
Plant survival dose-response of suspected protoporphyrinogen oxidase (PPO)-inhibitor-resistant (VRC) and PPO-sensitive (SIU and DSO) *Ambrosia trifida* populations to lactofen (A), fomesafen (B), glyphosate (C) and cloransulam-methyl (D) 21 days after application. Vertical lines indicate the herbicide rate estimated to reduce plant survival by 50% compared with the non-treated control group.

**Table 2.** Parameter estimates from log-logistic regression models (2-parameter log-logistic) describing plant survival responses of suspected protoporphyrinogen oxidase (PPO)-inhibitor-resistant (VRC) and PPO-sensitive (SIU and DSO) *Ambrosia trifida* populations to lactofen, fomesafen, glyphosate, and cloransulam-methyl 21 days after application.

| Herbicide | Population | Parameter estimates<sup>†</sup> |  |  |
| --- | --- | --- | --- | --- |
| | | b (± SE) | e (± SE) g a.i/a.e ha<sup>-1</sup> | Resistance ratio |
| Lactofen | VRC | 3.21 (1.41) | 87.05 (19.57) | - |
|  | DSO | 9.43 (36.03) | 82.53 (91.41) | 1.1 |
|  | SIU | 1.17 (0.24) | 23.45 (8.81) | 3.7 |
| Fomesafen | VRC | 1.57 (0.39) | 74.25 (23.63) | - |
|  | DSO | 3.96 (11.84) | 10.82 (9.05) | 6.9 |
|  | SIU | 7.25 (19.52) | 33.05 (124.15) | 2.2 |
| Glyphosate | VRC | 2.02 (0.78) | 7012.59 (2209.18) | 1.8 |
|  | DSO | 2.89 (0.80) | 3908.68 (543.88) | - |
|  | SIU | 2.29 (0.67) | 4340.69 (785.54) | 1.1 |
| Cloransulam-methyl | VRC | 0.84 (0.26) | 30.85 (9.89) | 6.9 |
|  | DSO | 0.67 (0.17) | 4.50 (2.05) | - |
|  | SIU | 0.56 (0.19) | 30.13 (14.20) | 6.7 |

<sup>†</sup> *b* is the slope around the effective rate, *e* denotes the effective herbicide rate necessary to induce 50% of plant mortality compared with the non-treated control group, SE is the standard error, and resistance ratio is calculated by dividing the estimated *e* value for the suspected resistant population (VRC) to that of the PPO-sensitive populations (SIU or DSO) from the fitted log-logistic models.

Overall, VRC exhibited reduced sensitivity to both PPO-inhibiting herbicides, with the clearest separation from the PPO-sensitive populations observed for fomesafen. Because of limited seed availability and the inability to collect additional seeds from the field where TMS was originally collected, only a single experimental run was conducted for this population (Supplementary Figures 1 and 2). For lactofen, the estimated LD_50_ values for TMS, VRC, and SIU, were 257.4 (±86.26), 165.41 (±73.9), and 0.29 (±1.1), respectively, indicating resistance ratios ranging from 887.7- to 570.3-fold. For fomesafen LD_50_ values for TMS, VRC, and SIU, were 318.40 (±111.94), 253.94 (±102.31) and 60.18 (±73.61), respectively, indicating resistance ratios ranging from 5.3- to 4.2-fold. Therefore, the TMS exhibited a higher resistance ratio to both PPO-inhibitors when compared to the VRC population.

**Figure 2.**
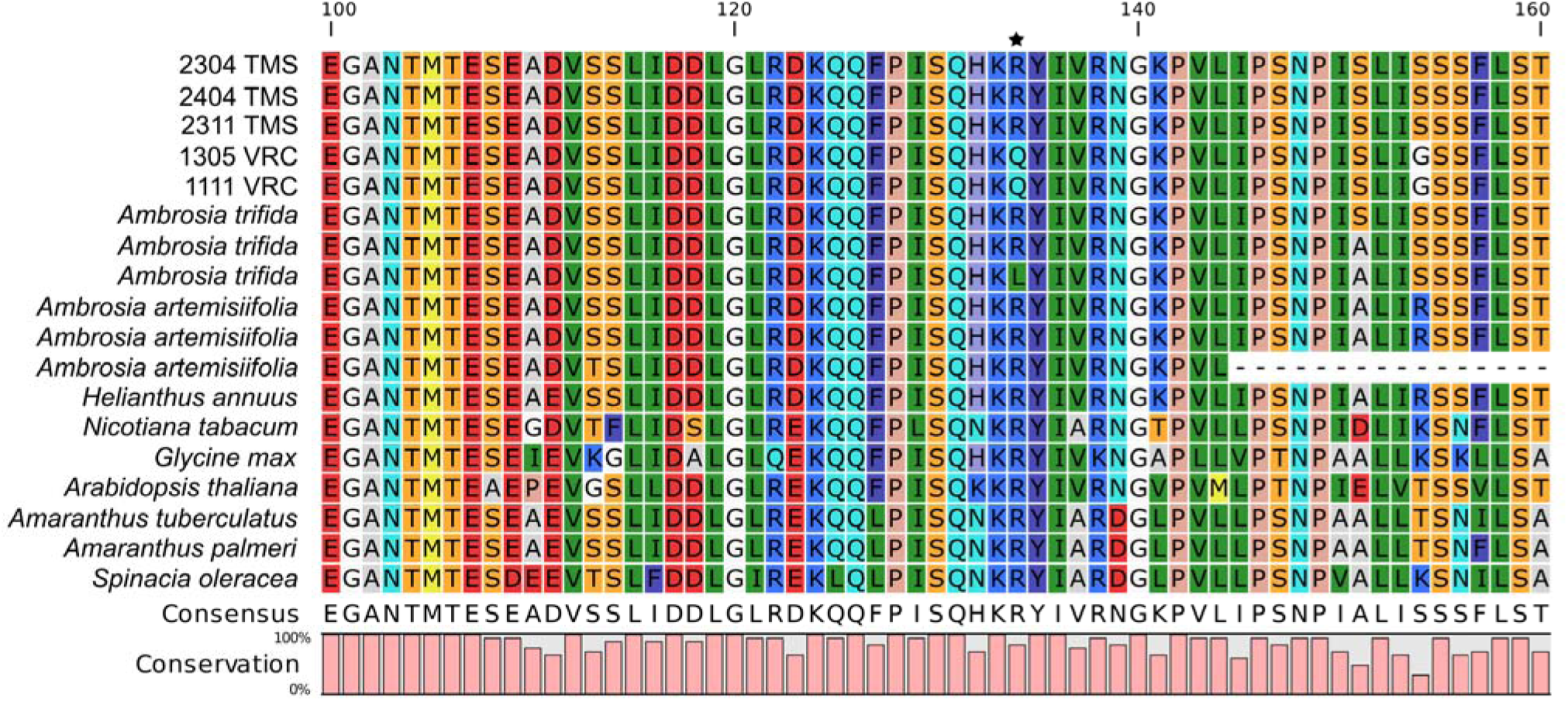
Multiple sequence alignment of PPO2 protein consensus sequences generated from suspected protoporphyrinogen oxidase (PPO)-resistant populations derived from the *Ambrosia trifida* TMS (2304, 2404, 2311) and VRC (1305, 1111) populations against publicly available PPO2 protein sequences of related and distant species. Amino acids are colored according to the RasMol color scheme indicating residues with similar biochemical properties, and the R98Q position suspected to contribute to PPO-inhibitor resistance is indicated by a star.

PPO-inhibiting herbicides have been an important tool for managing weeds in soybean fields in the midwestern and southern regions of the USA for more than 50 years (22). However, there was a notable decline in use at the turn of the century as farmers shifted to glyphosate as their primary option due to the introduction of glyphosate-resistant soybean in 1996 (43).

Nonetheless, with the current widespread glyphosate resistance in many troublesome weeds due to overreliance on this chemistry, PPO-inhibiting herbicide usage has increased as this site-of-action group offers an excellent alternative to control some glyphosate-resistant populations (22).

Glyphosate LD_50_ estimates for PPO-resistant and sensitive populations exceeded three times the labeled field rate of 1260 g a.e. ha^−1^. Estimated LD_50_ values were 7012.6 (±2209.2) g ai ha^-1^ for VRC, 4340.7 (±785.5) g ai ha^-1^ for SIU, and 3908.7 (±543.9) g ai ha^-1^ for DSO (Table 1; Figure 1C). These results indicate a substantial reduction in sensitivity to glyphosate across all three populations under greenhouse conditions. However, because SIU and DSO also showed high LD_50_ values, and no confirmed glyphosate-sensitive population was included, a glyphosate resistance ratio for VRC could not be reliably estimated. This response is consistent with the broader occurrence of glyphosate resistance in *A. trifida*, which has been reported in several Midwestern states, although it has not been confirmed in Illinois according to current resistance reports (12, 44, 45). Some of the mechanisms of resistance reported include reduced glyphosate translocation (46), rapid cell death response (47, 48), and the presence of a proline to serine substitution at amino acid position 106 of EPSPS, which is hypothesized to act additively or synergistically with other non-target-site (NTS) mechanisms to confer resistance in some phenotypes (49). Whether these mechanisms are present in VRC is unknown, although none of them exhibited the rapid cell death response.

For cloransulam-methyl, the DSO population was the most sensitive, with an estimated LD_50_ of 4.5 (±2.1) g a.i ha^-1^. In contrast, VRC and SIU had similar but higher LD_50_ values of 30.9 (±9.9) and 30.1 (±14.2) g a.i ha^-1^, respectively (Table 2; Figure 1D). These results indicate reduced sensitivity to cloransulam-methyl in VRC relative to DSO but not to SIU. Resistance to the ALS-inhibiting herbicides cloransulam-methyl and imazethapyr was first confirmed in *A. trifida* in Illinois in 1998 (12), sixteen years after the commercialization of the first ALS-inhibiting herbicide in 1982 (50). Resistance to ALS-inhibiting herbicides in *A. trifida* has been reported in other Midwestern states (12). One of the mechanisms of resistance identified in *A. trifida* is a tryptophan to leucine substitution at amino acid position 574 of ALS (13). When comparing the potential fitness between cloransulam-methyl-resistance and sensitive populations in the absence of herbicide, results showed equal competitive ability, fecundity, and seed viability (51).

The reduced sensitivity of VRC to PPO-, ALS-, and EPSPS-inhibiting herbicides is consistent with reports of PPO-inhibitor-resistant populations that also show resistance or reduced sensitivity to EPSPS- and/or ALS-inhibiting herbicides (52). For example, of the seven reported cases of PPO-inhibitor resistance in *A. artemisiifolia*, all populations also had evolved resistance to ALS-inhibitors, and three had evolved resistance to glyphosate (12). This is the same scenario observed in the VRC population, which exhibited reduced sensitivity to lactofen, fomesafen, glyphosate, and cloransulam-methyl. This is concerning because it reduces the number of effective postemergence herbicide options, especially for non-GMO soybeans, where reliance is concentrated on a limited set of herbicides (53, 54). Moreover, in fields where *A. trifida* populations already show reduced sensitivity to ALS- and EPSPS-inhibitors, PPO-inhibiting herbicides may become increasingly important, further increasing selection pressure on these herbicides (54).

### 3.2 Assessment of target-site mechanisms associated with resistance to PPO-inhibitors

A total of 18 polymorphisms were identified across all samples from the two populations in *PPX1*, of which 12 were synonymous and six were missense polymorphisms (Supplementary Table 3). In *PPX2*, 17 polymorphisms were identified, of which eight were synonymous and nine were missense polymorphisms (Supplementary Table 4). Of the missense substitutions identified in PPO1 and PPO2, all but one in PPO2, R98Q, occurred at positions that were not conserved (Figure 2; Supplementary Tables 5 and 6). Furthermore, R98Q was the only substitution located within the binding pocket of the catalytic domain, based on the crystal structure of tobacco PPO2 (PDB: 1SEZ; Supplementary Figure 3). This substitution was present in a heterozygous state in both biological replicates of the VRC population included in the transcriptome sequencing study but was absent from the PPO-sensitive population, SIU, and the suspected PPO-resistant population TMS (Figure 2).

We next evaluated whether *PPX1* or *PPX2* expression differed among the PPO-resistant populations VRC and TMS and the PPO-sensitive population SIU. No significant differences in *PPX1* or *PPX2* expression were detected among populations (Supplementary Figure 4). The absence of a known TS alteration in TMS suggests that PPO resistance in this population may involve a NTS mechanism of resistance. Although less documented than TS resistance, NTS resistance to PPO-inhibiting herbicides has been reported in several *Amaranthus* species, including *A. tuberculatus*, *A. palmeri*, and more recently *A. retroflexus* L. In these cases, enhanced herbicide metabolism has been implicated as an important mechanism, with cytochrome P450s, glutathione *S*-transferases, and UDP-glycosyltransferases among the gene families most frequently associated with resistance (55–59).

Substitutions at the PPO2 R98 site in *Ambrosia* spp. have previously been associated with resistance to PPO inhibitors. In *A. artemisiifolia*, an arginine-to-leucine substitution, R98L, was previously identified, and this same substitution was recently reported in a PPO-resistant *A. trifida* population from Wisconsin (11, 23). This arginine residue occurs within a highly conserved region of PPO2 across plant species and is positioned within the substrate-binding pocket of the catalytic domain, based on the crystal structure of tobacco PPO2 (39). However, residue numbering differs among taxa because of variation in the N-terminal region of the protein. For example, the homologous site is commonly referred to as R128 in Amaranthaceae species, where substitutions at this position have been implicated in resistance to PPO-inhibiting herbicides (60–62). Across PPO-resistant weed species, several substitutions at this conserved arginine residue have been reported, including arginine-to-leucine, -glycine, -methionine, and - isoleucine changes (R98L, R98G, R98M, and R98I, respectively) (52, 63).

Earlier research has shown that additional amino acid substitutions in PPO2 of *Ambrosia* spp. may also confer resistance to PPO-inhibiting herbicides. For example, previous *in vitro* and *in vivo* work using a transgenic *E. coli* cell line evaluated whether amino acid substitutions not yet reported in PPO-inhibitor resistance cases could arise in PPO2 and confer resistance. At the R98 site in *A. artemisiifolia* PPO2, R98G, R98L, R98P, R98Q, and R98W were predicted to be the substitutions most likely to confer resistance (64). However, prior to the present study, only R98L had been reported in PPO-inhibitor-resistant *Ambrosia* species, including *A. artemisiifolia* and *A. trifida* (11, 23). Of particular relevance to our findings, R98Q was among those substitutions shown to confer resistance to PPO-inhibiting herbicides in the *E. coli* system, which exhibited lactofen and fomesafen resistance ratios of approximately 5- and 15-fold relative to the wild type, respectively (64). The identification of R98Q in the VRC population suggests that this substitution likely contributes to the PPO-inhibitor resistance observed in this population, based on its localization at a conserved substrate-binding residue and functional evidence from *E. coli* studies.

With the results of this study, resistance to PPO-inhibiting herbicides has now been confirmed in at least three independent *A. trifida* populations across two states. Given the importance of these herbicides for managing *A. trifida* in non-GMO soybean and for controlling populations with reduced sensitivity or resistance to glyphosate, the recent occurrence of PPO-inhibitor resistance and its potential to spread is concerning. Therefore, integrated management strategies that combine chemical control with biological, cultural, and mechanical tactics, such as cover crops, tillage, electrical weed control, and harvest weed seed control, will be needed to reduce selection pressure and preserve the utility of PPO-inhibiting herbicides (65–69).

## 4 CONCLUSIONS

In conclusion, the suspected PPO-inhibitor-resistant populations VRC and TMS have evolved resistance to the PPO-inhibiting herbicides lactofen and fomesafen. VRC also exhibited reduced sensitivity to the ALS- and EPSPS-inhibiting herbicides cloransulam-methyl and glyphosate. However, because the reference populations SIU and DSO were not uniformly sensitive across all herbicide sites of action evaluated, further characterization using well-defined sensitive populations is needed to more accurately estimate the magnitude of reduced sensitivity to ALS- and EPSPS-inhibiting herbicides in VRC. Regarding PPO-inhibitor resistance, the R98L substitution previously reported in *A. artemisiifolia* and *A. trifida* was absent from *PPX2* in VRC; instead, VRC carried a different missense mutation at the same codon, resulting in an arginine-to-glutamine substitution, R98Q. This substitution was absent in TMS, and no significant differences in *PPX1* or *PPX2* expression were identified in either VRC or TMS. Together, these results suggest that R98Q is the most likely TS candidate associated with PPO-inhibitor resistance in VRC, whereas resistance in TMS may involve a NTS mechanism. To our knowledge, this represents the first report of the R98Q substitution associated with PPO-inhibitor resistance in plants.

## Author Contributions

C.B.R. collected seeds from field populations. C.B.R. and E.L. conducted the greenhouse studies. I.W.N., C.B.R., and E.L. performed the dose-response analyses. C.B.R. collected leaf tissue samples for the transcriptomic study. I.W.N. and A.J.L. conducted transcriptomic analyses. C.B.R. drafted the original manuscript. All authors reviewed and edited the manuscript. P.J.T. and K.L.G. acquired funding, conceptualized the study, and supervised the experiments.

## Acknowledgments

The authors thank the farmer for reporting the population (VRC) and providing the seeds, and the Southern Illinois University Carbondale weed science laboratory for technical assistance in the greenhouse. This research was partially funded by United Soybean Board (USB) Project #26-206-S-C-2-G.

## Data Availability Statement

The data that support the findings of this study are available from the corresponding author upon request.

